# Two distinct trophectoderm lineage stem cells from human pluripotent stem cells

**DOI:** 10.1101/762542

**Authors:** Adam Mischler, Victoria Karakis, Jessica Mahinthakumar, Celeste Carberry, Adriana San Miguel, Julia Rager, Rebecca Fry, Balaji M. Rao

## Abstract

Trophoblasts are the principal cell type of the placenta. The use of human trophoblast stem cells (hTSCs) as a model for studies of early placental development is hampered by limited genetic diversity of existing hTSC lines, and constraints on using human fetal tissue or embryos needed to generate additional cell lines. Here we report the derivation of two distinct stem cells of the trophectoderm lineage from human pluripotent stem cells. The first is a CDX2- stem cell equivalent to primary hTSCs – they both exhibit identical expression of key markers, are maintained in culture and differentiate under similar conditions, and share high transcriptome similarity. The second is a CDX2+ putative human trophectoderm stem cell (hTESC) with distinct cell culture requirements and differences in gene expression and differentiation relative to hTSCs. Derivation of hTSCs and hTESCs from pluripotent stem cells significantly enables construction of models for normal and pathological placental development.

## Introduction

Specification of the trophectoderm and the inner cell mass (ICM) is the first differentiation event during human embryonic development. The trophectoderm mediates blastocyst implantation in the uterus and is the precursor to all trophoblast cells in the placenta. Upon embryo implantation, the trophectoderm forms the cytotrophoblast (CTB), a putative stem cell that can differentiate to form the two major cell types in the placenta – the extravillous trophoblast (EVT) and the syncytiotrophoblast (STB) (Benirschke et al., 2012; Bischof and Irminger-Finger, 2005). The EVTs are involved in remodeling of uterine arteries, which is critical to ensure adequate perfusion of the placenta with maternal blood, whereas the multinucleated STB mediates the nutrient and gas exchange at the maternal-fetal interface (Moser et al., 2011; Yabe et al., 2016). Abnormalities in trophoblast development are associated with pregnancy-related pathologies such as miscarriage, preeclampsia and placenta accreta. Yet, despite its relevance to maternal and fetal health, constraints on research with human embryos and early fetal tissue impede mechanistic insight into early trophoblast development.

Trophoblast stem cells derived from first trimester human placental samples and blastocyst-stage embryos have emerged as an attractive in vitro model system for early human trophoblast (Okae et al., 2018). However, restricted accessibility of embryos and placental samples from early gestation and low genetic diversity of existing cell lines limit the use of this model. In contrast, pluripotent stem cells are a more accessible source for generation of in vitro models of human trophoblast. More importantly, unlike early gestation primary samples where the projected pregnancy outcome is uncertain, human induced pluripotent stem cells (hiPSCs) can potentially provide models of validated normal and pathological trophoblast development (Sheridan et al., 2019). However, whether *bona fide* trophoblast can be obtained from pluripotent stem cells has been a subject of intense debate (Roberts et al., 2014). A rigorous head-to-head comparison between trophoblast derived from pluripotent stem cells and their in vivo counterparts has proven difficult due to multiple reasons. Previous studies have used varying experimental protocols (Roberts et al., 2018), a self-renewing trophoblast stem cell population has not been derived from human pluripotent stem cells, and both primary placental samples and cultures of terminally differentiated trophoblast obtained from pluripotent stem cells exhibit heterogeneity and contain many cell types.

In this study, we report the derivation and maintenance of two distinct trophectoderm lineage stem cells from human embryonic stem cells (hESCs) and hiPSCs in chemically defined culture conditions. The first is a CDX2-human trophoblast stem cell (hTSC) that is comparable to primary hTSCs derived from early gestation placental samples. The second is a more primitive CDX2+ cell type that is a putative human trophectoderm stem cell (hTESC) (Knöfler et al., 2019). Critically, the isolation of self-renewing stem cell populations allowed a direct comparison of primary hTSCs with pluripotent stem cell derived hTSCs; genome wide transcriptomic analysis and functional differentiation assays establish their equivalence. The routine derivation of hTSCs and hTESCs from pluripotent stem cells will provide powerful tools for mechanistic studies on normal and pathological early trophoblast development.

## Results

### A chemically defined medium containing S1P enables differentiation of hESCs to CTB

Media formulations in previous studies on trophoblast differentiation of hESCs included components such as knockout serum replacement (KSR) or bovine serum albumin (BSA) that act as carriers for lipids. Pertinently, albumin-associated lipids have been implicated in activation of G-protein coupled receptor (GPCR)-mediated signaling (Mendelson et al., 2014; Yu et al., 2012). For instance, the phospholipid sphingosine-1 phosphate (S1P) present in KSR can activate YAP signaling; YAP plays a critical role in specification of the trophectoderm in mouse (Knott and Paul, 2014; Nishioka et al., 2008; Yagi et al., 2007). We investigated the use of S1P in the context of trophoblast differentiation of hESCs under chemically defined culture conditions, by modifying our previous protocol that utilized KSR (Sarkar et al., 2015, 2016). H1 and H9 hESCs cultured in E8 medium were differentiated for 6 days in E7 medium (E8 without TGFβ1) supplemented with S1P, by treatment with BMP4 and the activin/nodal inhibitor SB431542 (**Figure 1A**). Under these conditions, we observed upregulation of the CTB markers *CDX2* and *ELF5* (**Figure S1A, B**). Upregulation of *TBX4* was observed after 6 days. However, overall there were no significant changes in markers associated with neural or mesodermal differentiation after 6 days suggesting that differentiation to these lineages did not occur (**Figure S1A, B**). Immunofluorescence analysis at day 6 confirmed expression of the pan-trophoblast marker KRT7, and CTB markers P63 and GATA3 (**Figure 1B**; **Figure S1C**).

**Figure 1:**
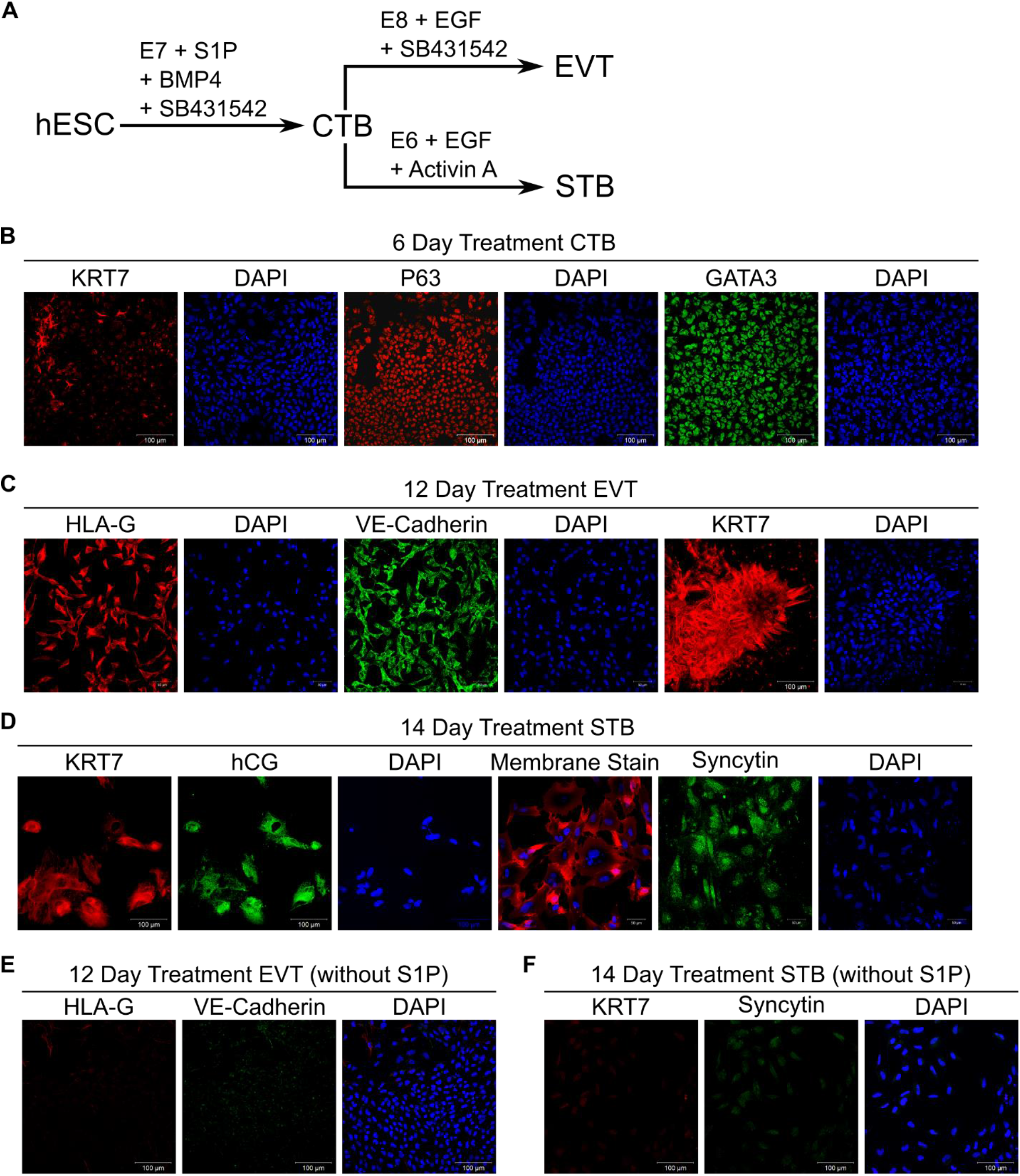
A chemically defined medium containing S1P enables differentiation of hESCs to CTB. (A) Schematic of protocol for hESC differentiation to trophoblast. (B) Immunostaining of KRT7, P63 and GATA3 in H9 hESCs at day 6 of initial treatment. Nuclei were stained with DAPI. (C) Confocal images of EVTs from 12-day treatment of H9 hESCs, staining for KRT7, HLA-G and VE-Cadherin. Nuclei were stained with DAPI. (D) Confocal images of STB from 14-day treatment of H9 hESCs, staining for KRT7, syncytin and hCG. Nuclei were stained with DAPI. (E) Confocal images of cells from 12-day EVT treatment of H9 hESCs upon removal of S1P, staining for HLA-G and VE-Cadherin. Nuclei were stained with DAPI. (F) Confocal images of cells from 14-day STB treatment of H9 hESCs upon removal of S1P staining for KRT7 and syncytin. Nuclei were stained with DAPI. Scale bars are 100μm for all images.

The putative CTB cells obtained at day 6 were investigated for their ability to differentiate to EVTs and STB, using protocols similar to those previously employed (Sarkar et al., 2015). Cells underwent an epithelial to mesenchymal transition over a 6-day period when passaged into E8 medium supplemented with epidermal growth factor (EGF) and SB431542. Immunofluorescence analysis showed expression of KRT7 and the EVT markers VE-Cadherin and HLA-G (**Figure 1C**, **S1D**). Alternatively, passaging CTB-like cells in E6 medium (E8 without TGFβ1 and bFGF) supplemented with activin and EGF resulted in formation of KRT7+ multinucleate cells expressing the STB markers hCG and syncytin over an 8-day period (**Figure 1D**, **S1E**). Removal of S1P from the medium during hESC differentiation to CTB-like cells abolished formation of EVTs that express HLA-G and VE-Cadherin (**Figure 1E**, **S2A)** under identical differentiation conditions (**Figure 1A**). Differentiation to STB also did not occur in the absence of S1P, as evidenced by lack of expression of syncytin and KRT7 (**Figure 1F**, **S2B**). Also, downregulation of the CTB marker *CDX2* and upregulation of transcripts of neural and mesoderm markers was observed in cells after 6 days of differentiation, upon removal of S1P (**Figure S2C**). Taken together these results show that CTB-like cells – similar to those in previous studies utilizing more complex culture conditions (Sarkar et al., 2015) – can be obtained by differentiation of hESCs in a chemically defined medium containing S1P. Further, inclusion of S1P is necessary for hESC differentiation to trophoblast in our chemically defined culture medium.

Rho GTPase signaling, downstream of GPCRs activated by S1P, has been implicated in nuclear localization of YAP (Mo et al., 2012; Ohgushi et al., 2015). Both Rho/RhoA associated kinase (ROCK) and nuclear YAP play a critical role in trophectoderm specification in the mouse (Kono et al., 2014; Nishioka et al., 2009). Therefore, we investigated the role of Rho/ROCK signaling and YAP in trophoblast differentiation of hESCs. The Rho/ROCK inhibitor Y-27632 was included during differentiation of hESCs to CTB-like cells and subsequent differentiation to EVT and STB to investigate the role of Rho/ROCK signaling. Under these conditions, HLA-G expression was observed in cells obtained from H9 hESCs; however, VE-Cadherin expression was weak and observed in only a few cells (**Figure S2D**). On the other hand, expression of EVT markers was not observed in cells derived from H1 hESCs. Additionally, presence of ROCK inhibition abolished STB formation, as shown by lack of expression of syncytin and KRT7 (**Figure S2E**).

To investigate the role of YAP signaling in trophoblast differentiation of hESCs, we used an hESC cell line (H9) that expresses an inducible shRNA against YAP (H9-YAP-ishRNA) or a scrambled shRNA control (Hsiao et al., 2016). YAP knockdown abolished differentiation to EVT and STB, as evidenced by lack of expression of the relevant markers. Notably, high cell death was observed (**Figure S2D, E**). Gene expression analysis revealed significant reduction in *ELF5* upon YAP knockdown, relative to the scrambled shRNA control (**Figure S2F**). Significant downregulation of the mesodermal genes *TBX4* and *LMO2* was observed, whereas *T* was upregulated, in H9-YAP-ishRNA, relative to the scrambled control. Taken together, these results show that Rho/ROCK signaling, and YAP are necessary for differentiation of hESCs to functional CTB that can give rise to both EVTs and STB, in our chemically defined culture medium.

### S1P mediates its effects on trophoblast differentiation of hESCs through its receptors

S1P acts through both receptor-mediated and receptor-independent pathways (Maceyka et al., 2012; Mendelson et al., 2014). To investigate the specific mechanism of S1P action during hESC differentiation to trophoblast, we replaced S1P with D-erythro-dihydrospingosine-1-phosphate (dhS1P) in our protocol. dhS1P acts as an agonist for the S1P receptors (S1PRs) but does not mediate an intracellular effect (Van Brocklyn et al., 1998). Replacing S1P with dhS1P yielded similar results – CTB-like cells showed high expression levels of CDX2, GATA3, P63, and TEAD4 (**Figure 2A**; **Figure S3A**). Upon further differentiation as previously described (**Figure 1A**), STB expressing KRT7 and hCG, and EVT expressing HLA-G and VE-Cadherin were obtained (**Figure 2B, C**; **Figure S3B, C**).

**Figure 2:**
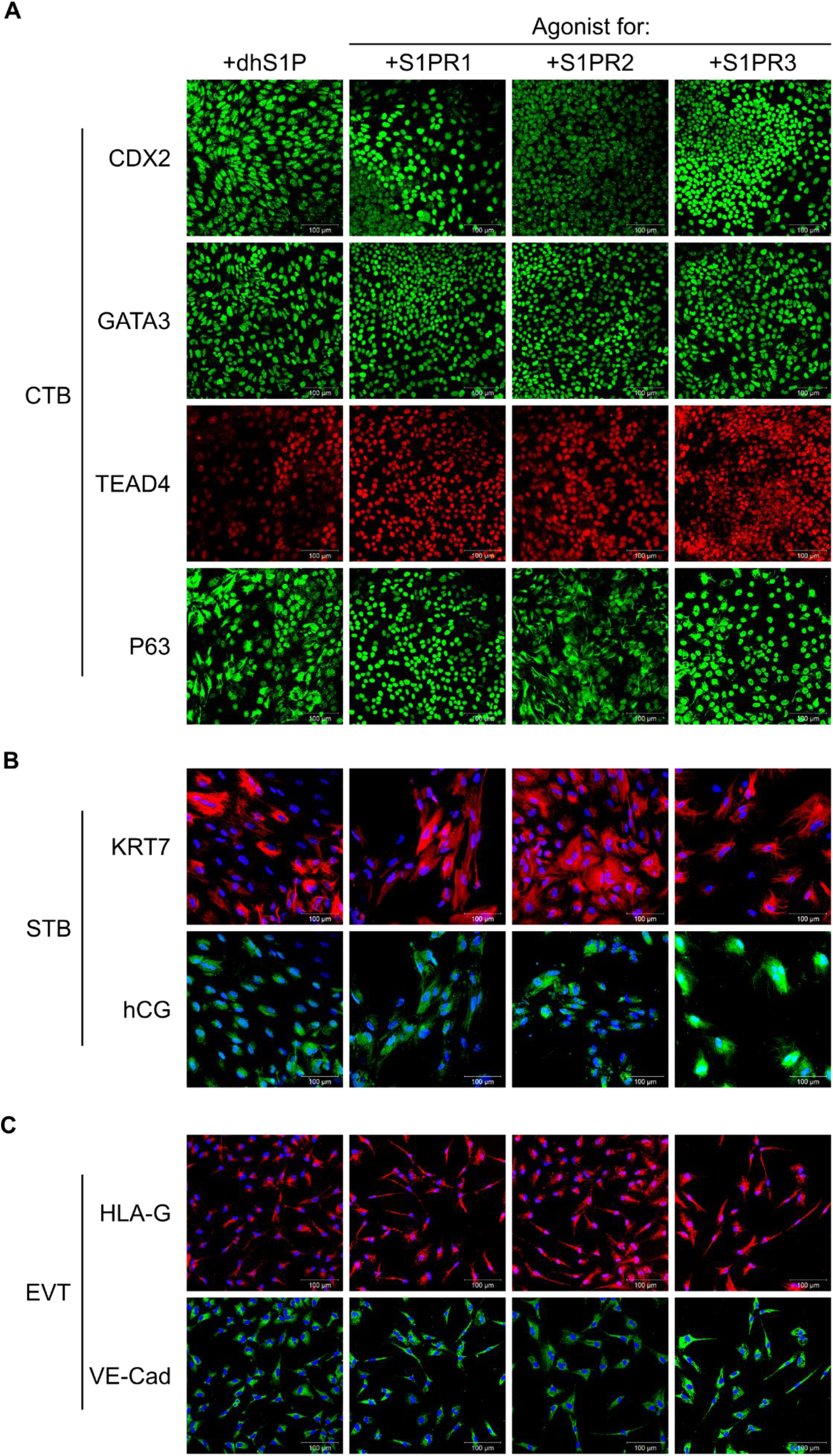
S1P mediates its effects on trophoblast differentiation of hESCs through its receptors. (A) Confocal images of CTB-like cells from 6-day treatment of H9 hESCs using D-erythro-dihydrospingosine-1-phosphate (dhS1P), CYM5442 (S1PR1 agonist), CYM5220 (S1PR2 agonist), and CYM5541 (S1PR3 agonist), staining for CDX2, GATA3, P63, and TEAD4. Nuclei were stained with DAPI. (B) Confocal images of STB cells from 14-day treatment of H9 hESCs using dhS1P, CYM5442, CYM5520, and CYM5541 during the initial 6-day treatment, staining for KRT7 and hCG. Nuclei were stained with DAPI. (C) Confocal images of EVT cells from 12-day treatment of H9 hESCs using dhS1P, CYM5442, CYM5220, and CYM5541 during the initial 6-day treatment, staining for HLA-G and VE-Cadherin. Nuclei were stained with DAPI. Scale bars are 100μm for all images.

S1P acts extracellularly through S1PR1-5 (Maceyka et al., 2012; Mendelson et al., 2014), however trophoblasts have been shown to only express S1PR1-3 (Johnstone et al., 2005). To identify specific S1PRs involved in trophoblast differentiation of hESCs in our culture system, we used selective chemical agonists for S1PR1-3 – CYM5442 hydrochloride, CYM5520 and CYM5541, respectively – to replace S1P in differentiation protocols previously discussed. Expression of CDX2, GATA3, P63, and TEAD4 was observed in CTB-like cells for all three agonists (**Figure 2A**; **Figure S3A**). Similarly, use of each agonist resulted in expression of the EVT markers HLA-G and VE-Cadherin, and formation of multinucleate STB expressing KRT7 and hCG (**Figure 2B, C**; **Figure S3B, C**). However, we observed some variability between the agonists. The intensity of CDX2 and P63 expression was higher with S1PR1 agonist CYM5442 and the S1PR3 agonist CYM5541. Nuclear P63 expression was strongest for CYM5442 compared to CYM5541. Notably, use of the S1PR2 agonist CYM5520 resulted in lower expression of CDX2, strong cytoplasmic expression of P63, and high heterogeneity in staining at day 6 relative to the other agonists. Formation of large multinucleated STB was more pronounced when the S1PR2 or S1PR3 agonists were used, as compared to the S1PR1 agonist. On the other hand, the S1PR1 and S1PR3 agonists enhanced formation of mesenchymal EVTs, relative to the S1PR2 agonist.

Taken together, our results show that receptor-mediated effects of exogenous S1P are sufficient for trophoblast differentiation of hESCs in our culture system. Since our qualitative observations showed that use of the S1PR3 agonist resulted in high CDX2 expression, and both multinucleate STB and mesenchymal EVTs could be obtained when the S1PR3 agonist was used, we chose the S1PR3 agonist for subsequent studies.

### Optimizing timing of hESC differentiation enables derivation of CDX2^+^ hTESCs

We investigated whether CTB-like cells obtained by treatment of hESCs with BMP4 and SB431542 in E7 medium supplemented with the S1PR3 agonist CYM5541 for 6 days could be passaged and maintained under conditions used for culture of primary hTSCs (Okae et al., 2018). Upon plating in trophoblast stem cell medium (TSCM) developed by Okae et al. (2018), hESC-derived CTB-like cells underwent differentiation, and epithelial colonies could not be retained after a single passage. CDX2 expression is upregulated significantly in as little as 2 days after initiation of hESC differentiation (**Figure S1A, B**). Additionally, previous studies have reported differentiation of hESCs to CDX2^+^/p63^+^ cells upon treatment with BMP for 4 days (Horii et al., 2016). Therefore, we explored the use of a shorter differentiation step for obtaining CTB-like cells. After 3 days of differentiation, H9 and H1 hESCs expressed nuclear CDX2, P63, and TEAD4 uniformly (**Figure 3B**). Quantitative image analysis showed that nearly all cells are CDX2+ at day 3, in contrast to CTB-like cells at day 6. Notably, use of a 6-day protocol resulted in significantly reduced fraction CDX2+ cells in the case of H1 hESCs in comparison to H9 hESCs; H9 cells retained CDX2+ cells longer (**Figure 3C**).

**Figure 3:**
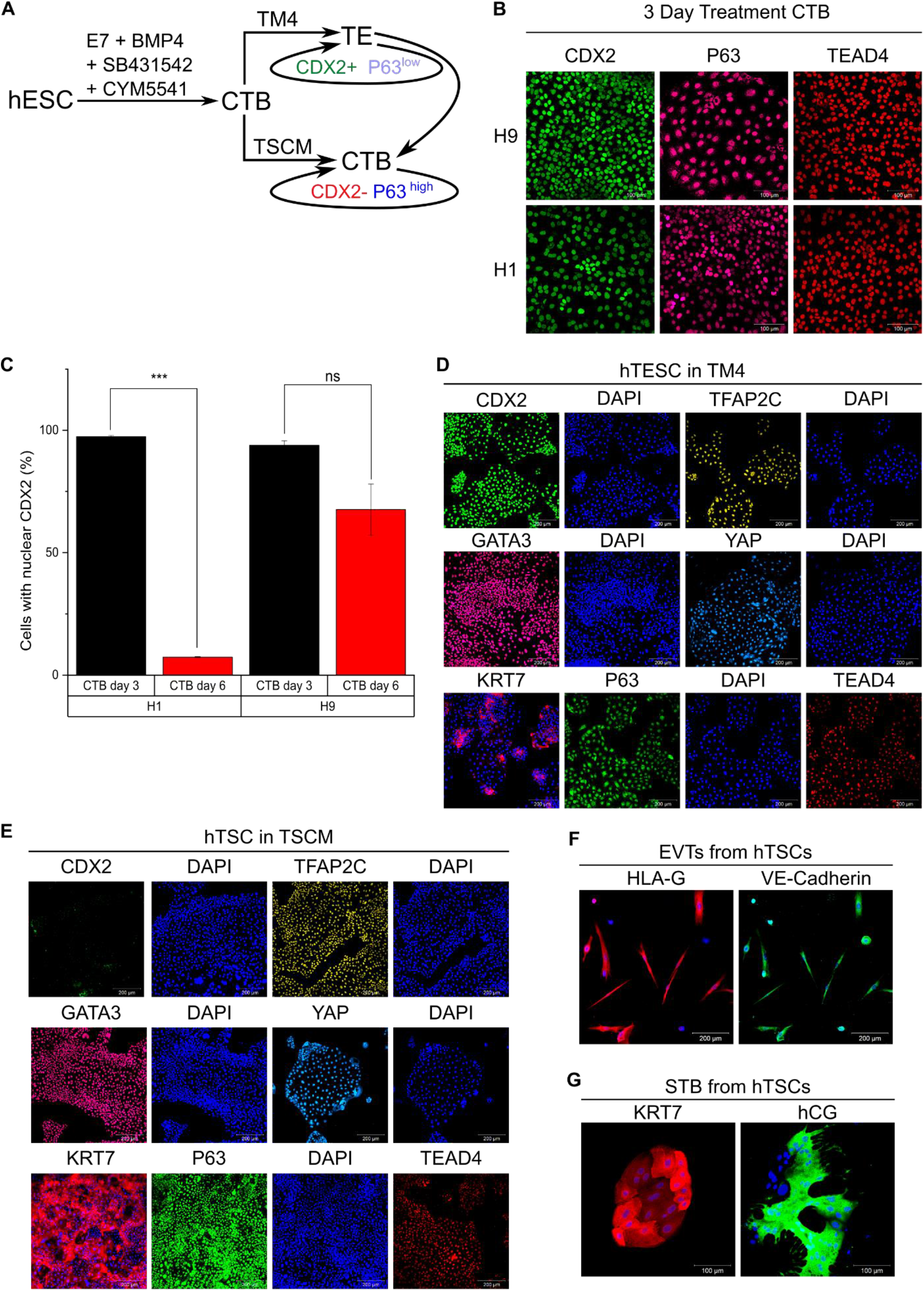
Optimizing timing of hESC differentiation enables derivation of CDX2+ hTESCs and P63+ hTSCs. (A) Schematic of differentiation protocol for establishment of hTESCs and hTSCs from hESCs. (B) Confocal images of 3-day treated H9 and H1 hESCs, staining for CDX2, P63, and TEAD4. Nuclei were stained with DAPI. Scale bars are 100μm. (C) Quantitative analysis of cells expressing nuclear CDX2 after 3-day and 6-day differentiation of H1 (day 3, n=5455; day 6, n=2448) and H9 (day 3, n=5552; day 6, n=6448) hESCs. Analysis was performed in MATLAB and at least 2 biological replicates were used. (Error bars are S.E., ***p<0.05). (D) Confocal images of H9 hESC-derived hTESCs in TM4, staining for CDX2, TFAP2C and GATA3, YAP, TEAD4, and P63. Nuclei were stained with DAPI. Scale bars are 200 μm. (E) Confocal images of H9 hESC-derived hTSCs in TSCM, staining for CDX2, TFAP2C and GATA3, YAP, TEAD4, and P63. Nuclei were stained with DAPI. Scale bars are 200 μm. (F) Confocal images of EVTs from H9 hESC-derived hTSCs, staining for HLA-G and VE-Cadherin. Nuclei were stained with DAPI. Scale bars are 200 μm. (G) Confocal images of STB from H9 hESC-derived hTSCs staining for hCG and KRT7. Nuclei were stained with DAPI. Scale bars are 100μm.

CDX2+ cells at day 3 were passaged into a chemically defined medium containing four major components (denoted TM4) – the S1PR3 agonist CYM5541, the GSK3β inhibitor CHIR99021, the TGFβ inhibitor A83-01, and FGF10. CHIR99021 and A83-01 are components of TSCM used for culture of primary hTSCs; FGF10 was included because FGFR2b signaling is active in primary hTSCs and the early placenta (Okae et al., 2018). Cells in TM4 could be maintained as epithelial colonies for 30+ passages over the course of 5 months. In TM4 medium, cells derived from H9 and H1 hESCs expressed the trophoblast markers CDX2, TFAP2C, YAP, TEAD4, and GATA3 (**Figure 3D**; **Figure S4A**) (Choi et al., 2012; Home et al., 2012; Nishioka et al., 2008; Niwa et al., 2005; Ralston et al., 2010; Yagi et al., 2007). Additionally, cells expressed the pan-trophoblast marker KRT7, and low levels of P63, which is expressed in CTBs found in the placental villi. Notably, CDX2 expression has been strongly associated with the trophectoderm and is lost once placental villi are formed (Blakeley et al., 2015; Hemberger et al., 2010; Horii et al., 2016; Knöfler et al., 2019). Due to their expression of CDX2, and to distinguish them from trophoblast stem cells that do not express CDX2, these cells are denoted as human trophectoderm stem cells (hTESCs).

### P63^+^ hTSCs derived from hESCs can be maintained in TSCM

We evaluated whether hTESCs could be maintained in TSCM used for culturing primary hTSCs (**Figure 3A**) (Okae et al., 2018). When hTESCs cultured in TM4 for 5+ passages were directly passaged into TSCM, cells underwent a change in colony morphology over ~ 3 passages; however, very little differentiation was observed. Notably, cell morphology of the hESC-derived cells closely resembled that of primary hTSCs in TSCM (**Figure S5A**) (Okae et al., 2018). Strikingly, however, hESC-derived hTESCs lost expression of CDX2 and gained high expression of P63. As discussed earlier, cells could be maintained as epithelial colonies when hESCs after 3 days of differentiation were passaged into TM4. In contrast, passaging day-3 differentiated hESCs into TSCM resulted in extensive differentiation, although a few epithelial colonies could be observed. Further passaging resulted in similar morphological changes in the epithelial colonies as those observed for hTESCs transitioning to TSCM. After ~ 6 passages, only epithelial colonies remained, and they closely resembled both the hTESCs transitioned into TSCM and primary hTSCs. H9 and H1 hESC-derived cells – passaged directly into TSCM after 3 days of differentiation or transitioned from TM4 (**Figure 3A**) – showed high expression of YAP, TEAD4, TFAP2C, and GATA3, similar to cells in TM4, but no expression of CDX2 (**Figure 3E**; **Figure S4B**). Lastly, hESC-derived hTSCs exhibit similar expression profile of trophoblast markers as primary hTSCs (**Figure S5B**). Therefore, these cells are denoted as hTSCs.

We further evaluated the differentiation potential of hESC-derived hTSCs using same protocols as those used by Okae et al. (2018) for differentiation of primary hTSCs to EVTs and STB (Okae et al., 2018). Similar to primary hTSCs, the hESC-derived hTSCs could be differentiated into mesenchymal EVTs expressing HLA-G and VE-Cadherin (**Figure 3F**; **Figure S4C**) and multinucleate cells expressing the STB markers hCG and KRT7 (**Figure 3G**; **Figure S4D**). Further, hESC-derived hTSCs retained their ability to differentiate into STB- and EVTs after 30 passages in TSCM. Strikingly however, hTESCs did not differentiate to EVTs using the same protocols used for hTSCs (**Figure S5C**). Taken together along with differences in culture conditions for maintenance and expression of the trophectoderm marker CDX2, these results suggest that hTESCs and hTSCs represent two distinct stem cell populations, with hTESCs being a more primitive cell type.

### Transcriptome analysis confirms equivalence of hESC-derived and primary hTSCs and reveals differences between hTSCs and hTESCs

We conducted genome wide transcriptome analysis on hTESCs, and hESC-derived and primary hTSCs using RNA sequencing. Principal component analysis (PCA) of transcriptomic signatures showed that hESC-derived and primary hTSCs cluster together, indicating similarities in overall gene expression (**Figure 4A**). Hierarchical clustering analysis further confirmed the very high transcriptome similarity between hESC-derived and primary hTSCs (**Figure 4B**). In conjunction with similarities in marker expression and culture conditions for maintenance and differentiation, these results establish the equivalence of hESC-derived and primary hTSCs.

**Figure 4:**
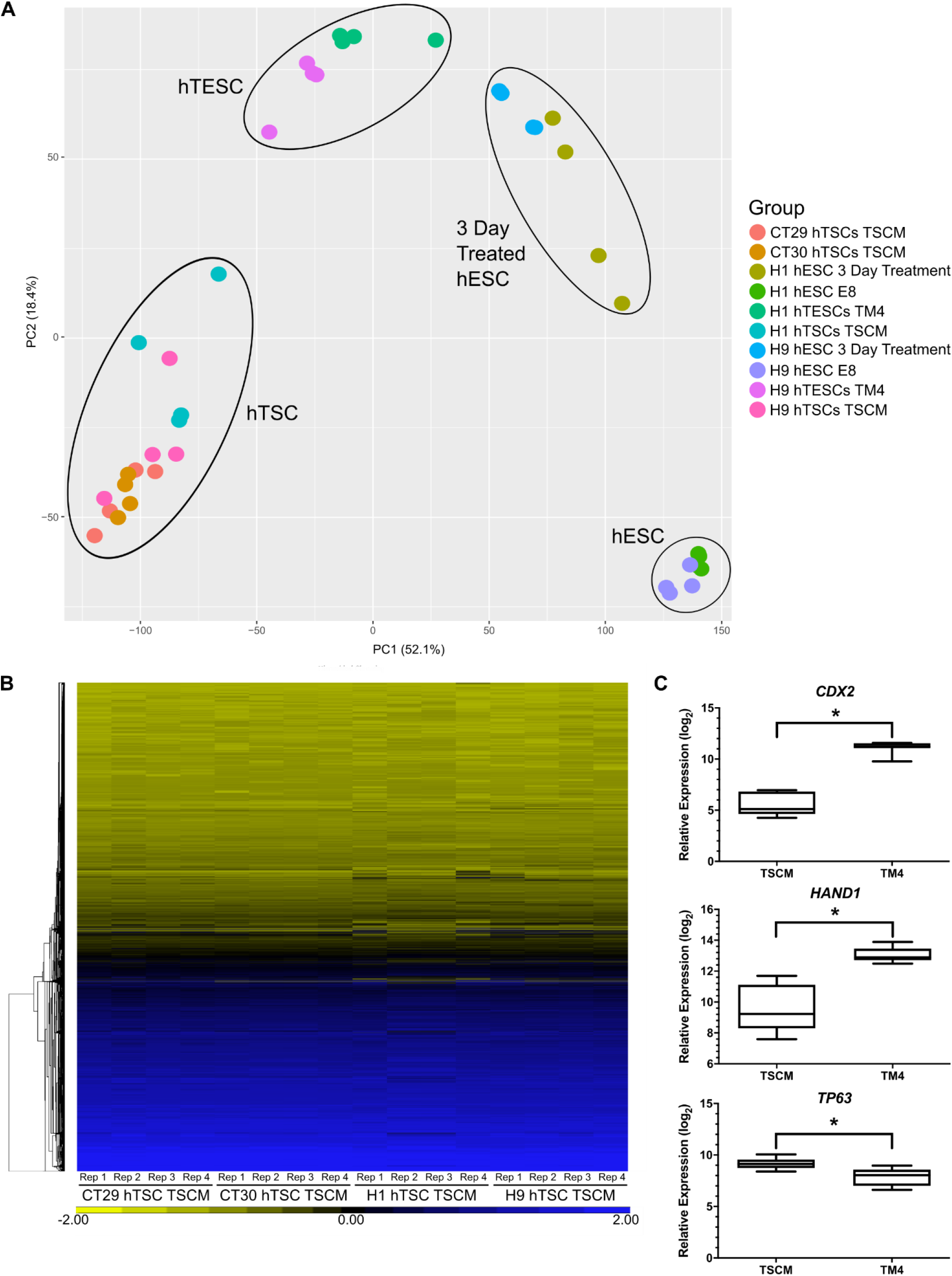
Transcriptome analysis confirms equivalence of hESC-derived and primary hTSCs and reveal differences between hTSCs and hTESCs. (A) Principal component analysis (PCA) of transcriptome data on H1 and H9 hESCs, H1 and H9 hESCs after 3-day treatment, H1 and H9 hESC-derived hTESCs cultured in TM4, H1 and H9 hESC-derived and primary (CT29 and CT30) hTSCs cultured in TSCM. (B) Hierarchical clustering analysis of transcriptome data from H1 and H9 hESC-derived and primary (CT29 and CT30) hTSCs. (C) Relative expression of trophectoderm-associated markers *CDX2* and *HAND1* and villous CTB-associated marker *TP63* in H1 and H9 hESC-derived hTESCs and hTSCs (*q<0.001).

PCA also showed that hTESCs are a distinct population of cells that cluster differently from hTSCs and hESCs differentiated to the trophoblast lineage for 3 days (**Figure 4A**). Higher expression of the trophectoderm-associated markers *CDX2* and *HAND1* is observed in hTESCs relative to hTSCs. On the other hand, expression of *TP63* – associated with villous CTB – is higher in hTSCs relative to hTESCs (**Figure 4C**). Statistical analysis of gene expression profiles identified genes that were significantly differentially expressed between hTESCs and hTSCs. Specifically, 269 genes showed significantly higher expression levels, and 275 genes showed significantly lower expression levels in hTESCs vs. hTSCs (**Tables S1 and S2**). Gene set enrichment analysis of these genes identified 300 and 47 gene ontology (GO) categories (out of 9996 queried categories) associated with genes showing higher and lower expression in hTESCs vs hTSCs, respectively (**Tables S3 and S4**). Interestingly, consistent with differences in colony morphology between hTESCs and hTSCs, genes associated with extracellular matrix, biological adhesion, and cell-cell adhesion were upregulated in hTESCs. Taken together along with distinct medium requirements for maintenance in cell culture, and differences in EVT differentiation under identical assay conditions, these results show that hTSCs and hTESCs represent distinct stem cell populations.

### HTESCs and hTSCs can be generated from hiPSCs

Lastly, we investigated if our results on derivation of hTESCs and hTSCs from hESCs can be extended to hiPSCs. Accordingly, we used our previously described protocols (**Figure 3A**) to derive hTESCs and hTSCs from the hiPSC line SBli006-A. HTESCs derived from SBli006-A hiPSCs maintained expression of CDX2, TFAP2C, GATA3, YAP KRT7, and TEAD4, along with low expression level of P63 in TM4 (**Figure 5A**). Similarly, hTSCs derived from SBli006-A hiPSCs expressed KRT7, P63, TEAD4, TFAP2C, YAP, and GATA3 in TSCM (**Figure 5B)**. Similar to the case with hESC-derived hTSCs, cells lost expression of CDX2 but gained higher expression levels of P63 and KRT7 in TSCM. Differentiation of hTSCs derived from SBli006-A hiPSCs using protocols described by Okae et al. (2018), resulted in formation of mesenchymal EVTs with high expression of HLA-G and VE-Cadherin (**Figure 5C**), and multinucleate STB expressing hCG and KRT7 (**Figure 5D**). These results show that hTESCs and hTSCs can also be derived from hiP-SCs.

**Figure 5:**
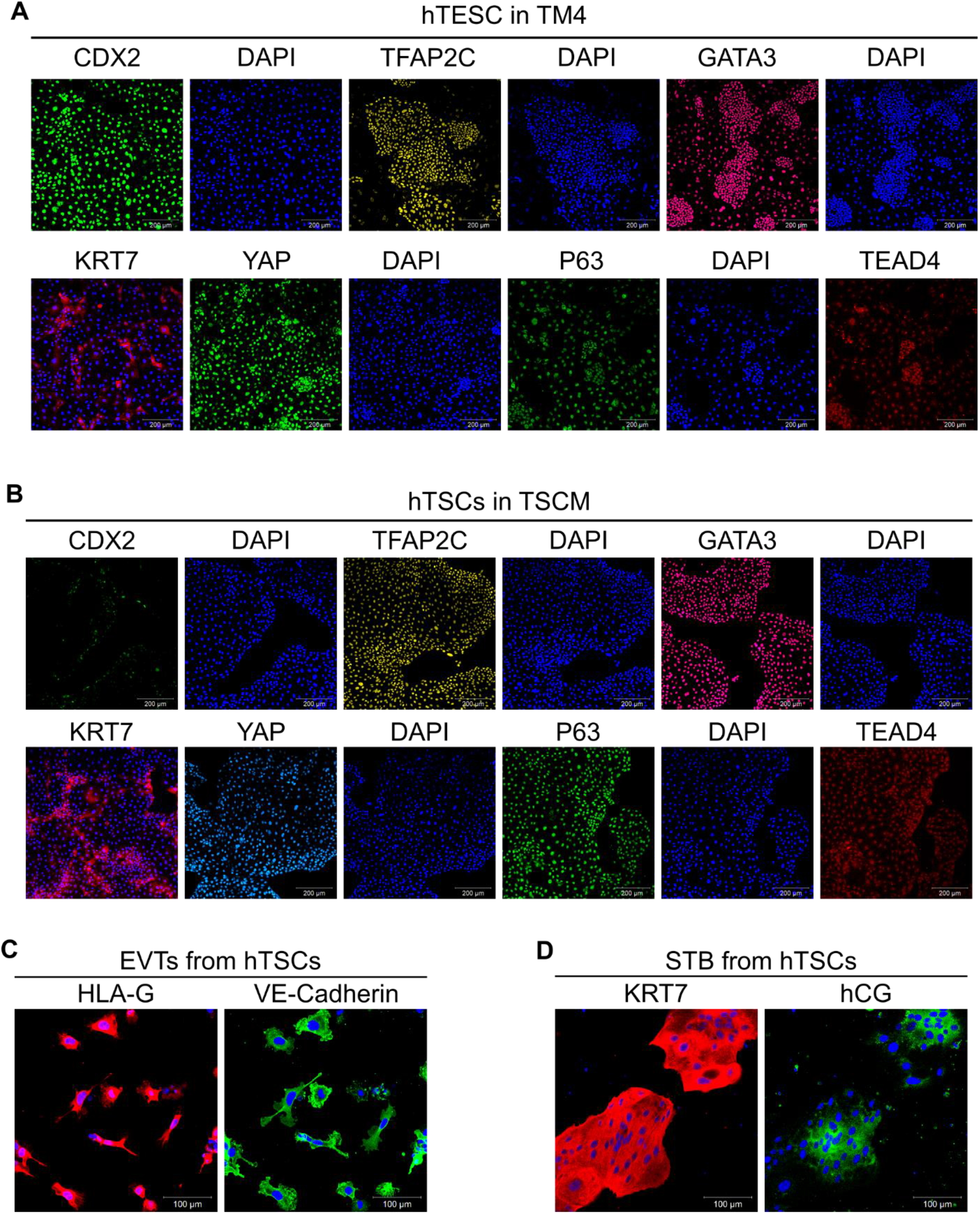
hTESCs and hTSCs generated from human iPSCs. (A) Confocal image of SBli006-A-derived hTESCs in TM4, staining for CDX2, TFAP2C, GATA3, YAP, TEAD4, and P63. Nuclei were stained with DAPI. Scale bars are 200μm. (B) Confocal images of SBli006-A-derived hTSCs in TSCM, staining for CDX2, TFAP2C, GATA3, YAP, TEAD4, and P63. Scale bars are 200μm. (C) Confocal images of EVTs from SBli006-A-derived hTSCs, staining for HLA-G and VE-Cadherin. Scale bars are 100μm. (D) Confocal images of STB from SBli006-A-derived hTSCs, staining for hCG and KRT7. Scale bars are 100μm.

## Discussion

In this study, we have shown that two distinct stem cell populations of the trophectoderm lineage – hTSCs and hTESCs – can be derived from hESCs and hiPSCs under chemically defined culture conditions. Whether bona fide trophoblast can be obtained from human pluripotent stem cells has been a subject of debate (Roberts et al., 2014). Despite extensive research in this area, conducting a rigorous head-to-head comparison between hESC-derived and primary trophoblasts has been challenging. The isolation of trophoblast stem cell populations from hESCs in this study, in conjunction with the recent derivation of primary hTSCs (Okae et al., 2018) enabled such a comparison. We have shown that hESCs can be differentiated to hTSCs that express markers consistent with primary hTSCs (P63, TEAD4, TFAP2C, YAP, and GATA3). The hESC-derived hTSCs are cultured in the same medium as primary hTSCs and differentiate to EVT and STB using similar protocols as those used for primary hTSCs. Further, hESC-derived hTSCs and primary hTSCs have highly similar transcriptomes. Taken together, these results establish the equivalence of hESC-derived and primary hTSCs and demonstrate that hESCs can indeed differentiate to bona fide trophoblasts.

### Role of receptor mediated S1P signaling and hESC culture medium in trophoblast differentiation of hESCs

Previous studies on trophoblast differentiation of hESCs have employed differing protocols, resulting in significantly different outcomes in some cases. Notably, Bernardo et al. (2011) reported that BMP treatment of hESCs results in differentiation of hESCs to mesoderm and not trophoblast. Our studies suggest two potential explanations for discrepancies in previous studies. First, our results show that receptor-mediated signaling by the albumin-associated sphingolipid S1P plays a critical role in hESC differentiation to trophoblast in our medium. Differences in results reported by previous studies may be due to variability in the lipid composition of media used during trophoblast differentiation of hESCs. Second, the medium used for routine maintenance of undifferentiated hESCs likely contributes significantly to their differentiation potential. For instance, unlike hESCs cultured in the presence of KSR, hESCs in E8 medium exhibit features of naïve pluripotency (Cornacchia et al., 2019). Recent studies report the conversion of hESCs to expanded potential stem cells (EPSCs) by transitioning hESCs to a human EPSC medium. Significantly, hTSC-like cells can be obtained by passaging EPSCs – but not hESCs in KSR containing medium – in TSCM used for culture of primary hTSCs (Gao et al., 2019). The efficiency of establishing hTSC-like lines was low (~ 30% with manual isolation of colonies) and a rigorous transcriptome comparison with primary hTSCs was not conducted. Nevertheless, taken together these studies underscore the importance of hESC culture conditions in their differentiation potential. Differences in culture conditions for undifferentiated hESCs may lead to inconsistent results during trophoblast differentiation of hESCs.

Our results show that Rho/ROCK signaling and YAP are necessary for trophoblast differentiation of hESCs in our medium. However, the exact molecular mechanisms that underlie acquisition of trophoblast fate in the presence of S1P receptor activation need to be further studied. Our results do not preclude the possibility that Rho/ROCK and/or YAP acts independently of S1P receptor mediated signaling in our system.

### Differences between hTESCs and hTSCs

Marker expression analysis, functional differentiation assays, and genome-wide transcriptome analysis confirm the equivalence of hESC-derived hTSCs and primary hTSCs that are similar to villous CTB. However, hTESCs differ significantly from hTSCs. They do not undergo differentiation to EVTs under the culture conditions used for differentiating hESC-derived and primary hTSCs. Transcriptome analysis shows that genes associated with several key pathways and biological processes are differentially regulated between hTESCs and hTSCs. Significantly, hTESCs – but not hTSCs – express high levels of the trophectoderm-associated markers *CDX2* and *HAND1*. Therefore we opt for the nomenclature human trophectoderm stem cells (hTESCs) as proposed by Knöfler et al. (2019), to distinguish these CDX2+ stem cells from hTSCs. Consistent with the more primitive nature of hTESCs, they can be readily transitioned into TSCM used for culturing hTSCs, as was seen by Okae et al. (2018) when transitioning trophectoderm cells of blastocysts into TSCM. Subsequently, similar to primary hTSCs, hTESCs lose expression of CDX2 and express higher levels of P63 in TSCM, and can differentiate to form EVTs and STB. Note that primary hTSCs derived from the trophectoderm in the blastocyst stage embryo lose expression of CDX2 in TSCM (Okae et al., 2018). On the other hand, it has not been possible yet to revert hTSCs to hTESCs by culturing in TM4 medium. Further studies are needed to rule out the possibility of such a reversion to the more primitive cell type.

### Considerations for derivation and culture of hTESCs

To derive hTESCs, undifferentiated hESCs maintained in E8 medium are first treated for 3 days with the S1PR3 agonist, BMP4 and the activin/nodal inhibitor SB4315432, to obtain CDX2+ cells. Subsequently, CDX2+ cells are passaged in TM4 medium to obtain hTESCs. Using this protocol, we observed increased differentiation of H1-derived cells upon passage into TM4 medium, relative to H9- and SBli006-A-derived cells. Shortening the initial treatment step in case of H1 hESCs to 2 days eliminated excessive differentiation and facilitated derivation of hTESCs. However, we were unable to derive hTESCs with any cell line when the initial treatment was greater than 3 days.

In our studies, the initial hESC differentiation step was carried out in E7 medium that contains bFGF. Differentiation of hESCs to trophoblast has been carried out in the presence or absence of exogenous FGF (Amita et al., 2013; Das et al., 2007). Consistent with this, we found that hTESCs could be formed even if the initial treatment was carried out in E6 medium lacking bFGF, instead of E7 medium.

It is important to note that hTESCs proliferate slower in culture than hTSCs. We also observe that the attachment of hTESCs to tissue culture plates is less efficient than hTSCs. Finally, we observe that excessive differentiation in TM4 medium during early passages could be countered by reducing the concentration of ascorbic acid (32 μg/mL instead of 64 μg/mL) in TM4. Additional studies on composition of TM4 medium or the substrates used to coat tissue culture plates may lead to improved growth rate and attachment efficiency. Alternatively, the slower growth rate and less efficient attachment characteristics may be an inherent feature of the hTESC state. Nonetheless, we have successfully maintained hTESCs derived from all cell lines studied for at least 20 passages, in several independent runs over 5+ months. We recommend passaging hTESCs routinely at higher cell densities relative to hTSCs, and troubleshooting cell line specific variability by optimizing the initial treatment step and/or lowering ascorbic acid concentration in TM4.

### Derivation of hTSCs from hiPSCs

We have shown that hTESCs and hTSCs can be derived from hiPSCs. Since hiPSCs can be derived by reprogramming easily accessible somatic tissues, hTSCs and hTESCs derived from hiPSCs can greatly accelerate research in placental biology. Further, arguably a limitation of primary hTSCs is that pregnancy outcomes at term for the early gestation placental samples or blastocyst stage embryos used cannot be predicted accurately. In contrast, hiPSC-derived hTSCs and hTESCs, from hiPSCs generated using somatic tissues obtained at term, will enable development of models of validated normal and pathological trophoblast development. Pertinently, Sheridan et al. (2019) have derived hiPSCs from umbilical cords of normal pregnancies and those associated with early onset preeclampsia. Our results also gain particular significance in the light of restrictions on research with fetal tissue (Du Toit, 2019).

In conclusion, using optimized cell culture protocols detailed in the current study, we have derived two distinct stem cell populations of the trophectoderm lineage – hTESCs and hTSCs – from human pluripotent stem cells. These stem cell models will be powerful tools for in vitro studies on human trophoblast development.

## Materials and Methods

### Key resources table

| REAGENT or RESOURCE | SOURCE | IDENTIFIER |
| --- | --- | --- |
| **Antibodies** |  |  |
| Anti-KRT7 | Santa Cruz Biotechnology | Cat#sc-23876, RRID:AB_2265604 |
| Anti-KRT7 | Cell Signaling Technologies | Cat# 4465, RRID:AB_11178382 |
| Anti-hCG | Abcam | Cat# ab9582, RRID:AB_296507 |
| Anti-hCG | Abcam | Cat# ab9376, RRID:AB_307221 |
| Anti-P63 | Cell Signaling Technologies | Cat# 13109, RRID:AB_2637091 |
| Anti-GATA3 | Cell Signaling Technologies | Cat# 5852, RRID:AB_10835690 |
| Anti-TFAP2C | Cell Signaling Technologies | Cat# 2320, RRID:AB_2202287 |
| Anti-YAP | Cell Signaling Technologies | Cat# 4912, RRID:AB_2218911 |
| Anti-TEAD4 | Abcam | Cat# ab58310, RRID:AB_945789 |
| Anti-CDX2 | Abcam | Cat# ab76541, RRID:AB_1523334 |
| Anti-VE-Cadherin | Cell Signaling Technologies | Cat# 2500, RRID:AB_10839118 |
| Anti-HLA-G | Abcam | Cat# ab52455, RRID:AB_880552 |
| Anti-Syncytin | Santa Cruz Biotechnology | Cat# sc-50369, RRID:AB_2101536 |
| Rabbit Polyclonal IgG | R&D Systems | Cat# AB-105-C, RRID:AB_354266 |
| Rabbit XP IgG | Cell Signaling Technologies | Cat# 3900, RRID:AB_1550038 |
| Mouse IgG1 | Abcam | Cat# ab18447, RRID:AB_2722536 |
| Mouse IgG2a | Abcam | Cat# 554126, RRID:AB_479661 |
| Alexa Fluor 488-conjugated anti-rabbit IgG | Thermo Fisher Scientific | Cat# A-11034, RRID:AB_2576217 |
| Alexa Fluor 647-conjugated anti-rabbit IgG | Thermo Fisher Scientific | Cat# A-21052, RRID:AB_2535719 |
| DAPI | R&D Systems | Cat#5748 |
| **Chemicals, Peptides, and Recombinant Proteins** |  |  |
| TrypLE | Thermo Fisher Scientific | Cat#12604013 |
| Vitronectin | Thermo Fisher Scientific | Cat#A14700 |
| Laminin 521 | Stem Cell Technologies | Cat#77003 |
| TeSR-E8 | Stem Cell Technologies | Cat#05990 |
| TeSR-E7 | Stem Cell Technologies | Cat#05914 |
| TeSR-E6 | Stem Cell Technologies | Cat#05946 |
| ReLeSR | Stem Cell Technologies | Cat#05872 |
| Sphingosine-1-phosphate | Tocris | Cat#1370 |
| D-erythro-dihydrosphingosine-1-phosphate | Abcam | Cat#ab141750 |
| SB431542 | Tocris | Cat#1614 |
| BMP4 | Thermo Fisher Scientific | Cat#PHC9534 |
| CYM5442 hydrochloride | Tocris | Cat#3601 |
| CYM5520 | Tocris | Cat#5418 |
| CYM5541 | Tocris | Cat#4897 |
| Y-27632 dihydrochloride | Tocris | Cat#1254 |
| EGF | R&D Systems | Cat#236-EG |
| Doxycycline hyclate | Tocris | Cat#4090 |
| Puromycin dihydrochloride | Tocris | Cat#4089 |
| Activin A | R&D Systems | Cat#338-AC |
| Greiner Bio-one Cell View glass plates | Greiner Bio-one | Cat#627965 |
| 4% Paraformaldehyde in PBS | Thermo Fisher Scientific | Cat#R37814 |
| Triton X-100 | Sigma | Cat#T8787 |
| PBS w/o CaMg | Sigma | Cat#D5773 |
| PBS w/ CaMg | Sigma | Cat#D8662 |
| Human IgG | Immunoreagents | Cat#Hu-003-C |
| BSA | Fisher Scientific | Cat#BP9703 |
| 10% BSA fatty acid free in PBS | Sigma | Cat#A1595 |
| VPA | Sigma | Cat#P6273 |
| A83-01 | Tocris | Cat#2939 |
| 2-mercaptoethanol | Sigma | Cat#M3148 |
| FBS | Thermo Fisher Scientific | Cat#16141-061 |
| DMEM/F12 | Thermo Fisher Scientific | Cat#11320033 |
| ITS-X | Thermo Fisher Scientific | Cat#51500-056 |
| L-ascorbic acid | Sigma | Cat#A8960 |
| Pen/Strep | Thermo Fisher Scientific | Cat#15140122 |
| Forskolin | Tocris | Cat#1099 |
| Neuregulin | Cell Signaling Technologies | Cat#5218SC |
| Matrigel | Corning | Cat#354234 |
| KSR | Thermo Fisher Scientific | Cat#10828028 |
| Trizol Reagent | Thermo Fisher Scientific | Cat#15596018 |
| DEPC | Sigma | Cat#95284 |
| Baseline Zero DNAase Kit | VWR | Cat#76081-624 |
| Oligo-dT | IDT | Cat#51-01-15-07 |
| dNTP mix | Thermo Fisher Scientific | Cat#10297018 |
| Superscript II RT | Thermo Fisher Scientific | Cat#18064014 |
| SYBR Green Supermix | Bio-rad | Cat#1725272 |
| Methanol | Fisher Scientific | Cat#A412-500 |
| Acetone | Fisher Scientific | Cat#A18-500 |
| **Critical Commercial Kits** |  |  |
| GeneJET RNA Purification Kit | Thermo Fisher Scientific | Cat#K0731 |
| **Oligonucleotides** |  |  |
| qPCR Primers | IDT | Methods S1 for primer sequences |
| **Software and Algorithms** |  |  |
| R (v3.6.0) | http://www.R-project.org/ | N/A |
| DESeq2 package (v1.22.2) |  |  |
| SAS Software |  | N/A |
| Zeiss Zen Software | https://www.zeiss.com/microscopy/us/products/microscope-software/zen-lite.html | N/A |

### Culture of hESCs and hiPSCs

H1 and H9 hESCs and SBli006-A hiPSCs were cultured on plates coated with vitronectin (5 μg/ml) at room temperature for at least one hour. Cells were cultured in 2 ml of TeSR-E8 medium at 37°C in 5% CO_2_ in 6-well plates and culture medium was replaced every day. When cells reached confluency, they were passaged using ReLeSR according to the manufacturer’s protocol, at a 1:10 split ratio.

### Differentiation of hESCs (6 days protocol)

The day after passaging, differentiation was initiated in H1 or H9 hESCs by treatment with S1P (10 μM), SB431542 (25 μM) and BMP4 (20 ng/ml) in TeSR-E7 for 6 days. In some experiments, the S1PR agonists CYM5442 hydrochloride (10 nM), CYM5520 (5 μM), CYM5541 (2 μM), or the Rho/ROCK inhibitor Y-27632 (5 μM), or doxycycline (2 μM), and/or puromycin (1.5 μg/mL) was added during the differentiation process. The medium was replaced every day. At day 6 of treatment, cells were dissociated with TrypLE for 5 min at 37°C. For differentiation to EVTs, cells were seeded in a 6-well plate pre-coated with 5 μg/ml of vitronectin at a density of 7×10^4^ cells per well and cultured in 2 ml of TeSR-E8 medium supplemented with SB431542 (25μM) and EGF (2.5 ng/ml). Medium was replaced every other day and analyzed at day 12 of total treatment. For differentiation to STB, cells were seeded in a 6-well plate pre-coated with 5 μg/ml of vitronectin at a density of 4×10^4^ cells per well and cultured in 2 ml of TeSR-E6 supplemented with Activin A (20 ng/ml) and EGF (50 ng/ml). Medium was replaced every other day and analyzed at day 14 of total treatment.

To investigate the role of YAP signaling in TB formation from hESCs, we used an hESC cell line (H9) that expresses an inducible shRNA against YAP (H9-YAP-ishRNA) (Hsiao et al., 2016). This cell line along with a scrambled shRNA control were a kind gift from Dr. Sean Palecek (University of Wisconsin). shRNA expression was induced with doxycycline under constant exposure to puromycin as the selection marker.

### Differentiation of hESCs to hTESCs and hTSCs

The day after passaging, hESCs were differentiated by treatment with CYM5541 (2 μM), SB431542 (25 μM), BMP4 (20 ng/ml) in TeSR-E7 for 2 and 3 days for H1 and H9 hESCs, respectively. The medium was replaced every day. After 2 or 3 days of treatment, cells were dissociated with TrypLE for 5 minutes at 37°C. For propagation of hTESCs, all cells were seeded in a 6-well plate pre-coated with 3 μg/ml of vitronectin and 1 μg/ml of Laminin 521 at a density of ~5×10^4^ cells per well and cultured in 2 ml of TM4 medium [TeSR-E6 medium supplemented with CYM5541 (2 μM), A 83-01 (0.5 μM), FGF10 (25ng/ml) and CHIR99021 (2 μM)]. For establishment of hTSCs, all cells were seeded in a 6-well plate pre-coated with 3 μg/ml of vitronectin and 1 μg/ml of Laminin 521 at a density of ~5×10^4^ cells per well and cultured in 2 ml of TSCM developed by Okae et al. (2018) [DMEM/F12 supplemented with 0.1 mM 2-mercaptoethanol, 0.2% FBS, 0.5% Penicillin-Streptomycin, 0.3% BSA, 1% ITS-X supplement, 1.5 μg/ml L-ascorbic acid, 50 ng/ml EGF, 2 μM CHIR99021, 0.5 μM A83-01, 1 μM SB431542, 0.8 mM VPA and 5 μM Y27632]. HTESCs were directly passaged into TSCM for formation of hTSCs; complete transition took ~ 5 passages. Alternatively, hESC after 2 or 3 days of differentiation were directly passaged into TSCM.

### Culture of hTESCs and hTSCs

HTESCs and hTSCs were cultured in TM4 and TSCM, respectively, in 2 ml of culture medium at 37C in 5% CO_2_. Culture medium was replaced every 2 days. When hTESCs or hTSCs reached 70-90% confluence, they were dissociated with TrypLE at 37°C for 5-10 minutes and passaged to a new 6-well plate pre-coated with 3 μg/ml of vitronectin and 1 μg/ml of Laminin 521 at a 1:3-1:4 split ratio for hTESCs and 1:4-1:6 split ratio for hTSCs. hTESCs grown in TM4 medium were supplemented with Y-27632 upon passage to aid in single cell attachment. Cells were routinely passaged approximately every 4-6 days. hTESCs and hTSCs at passages 5+ were used for analysis, with the exception of one replicate of H1-derived hTESCs used in RNA-sequencing analysis where cells at passage 2 in TM4 were used.

CT29 and CT30 primary hTSCs were a kind gift from Dr. Hiroaki Okae (Tohoku University, Japan (Okae et al., 2018)). Primary hTSCs were cultured in TSCM, similar to hESC-derived hTSCs.

### Differentiation of hTESCs and hTSCs

hTSCs were grown to ~80-90% confluence in TSCM and dissociated with TrypLE for 10 min at 37°C. For differentiation to EVTs and STB, slightly modified versions of protocols developed by Okae et al. (2018) were used. For differentiation to EVTs, hTSCs were seeded in 6-well plates pre-coated with 3 μg/ml vitronectin and 1 μg/ml of Laminin 521 at a density of 1.25×10^5^ cells per well and cultured in 2 mL of EVT medium (DMEM/F12 supplemented with 0.1 mM 2-mercaptoethanol, 0.5% Penicillin-Streptomycin, 0.3% BSA, 1% ITS-X supplement, 100 ng/ml NRG1, 7.5 μM A83-01, 2.5 μM Y27632, and 4% KSR). Matrigel was added to a final media concentration of 2% after suspending the cells in EVT medium. At day 3, the medium was replaced with the EVT medium without NRG1, and Matrigel was added to a final concentration of 0.5%. At day 6, cells were dissociated with TrypLE for 15 min at 37°C and passaged to a new vitronectin/laminin-coated 6-well plates at a 1:2 split ratio. The cells were suspended in the EVT medium without NRG1 and KSR. Matrigel was added to a final concentration of 0.5%, and cells were analyzed after two additional days of culturing. For differentiation to STB, cells were seeded in 6-well plates pre-coated 3 μg/ml vitronectin and 1 μg/ml of Laminin 521 at a density of 1.5×10^5^ cells per well and cultured in 2 mL of DMEM/F12 supplemented with 0.1 mM 2-mercaptoethanol, 0.5% Penicillin-Streptomycin, 0.3% BSA, 1% ITS-X supplement, 2.5 μM Y27632, 2 μM forskolin, and 4% KSR. The medium was replaced at day 3, and cells were analyzed at day 6.

### RNA Isolation, cDNA synthesis and Quantitative PCR

RNA was isolated using Trizol™ reagent using the manufacturer’s protocol. For cDNA synthesis, the RNA pellet was dissolved in diethyl pyrocarbonate (DEPC)-treated water. The RNA was purified using Baseline-ZERO DNase buffer and Baseline-ZERO DNase enzyme and incubating at 37°C for 30 min. The purification was stopped with Baseline-ZERO DNase stop solution and heated at 65°C for 10 min. cDNA was synthesized using 18-mer Oligo-dT and dNTP mix and heated to 65°C for 5 min and quickly chilled on ice. First strand buffer and DTT was added and incubated at 42°C for 2 min then superscript II RT enzyme was added and incubated at 42°C for 50 min. The enzyme was inactivated at 70°C for 15 min. The cDNA was stored at −20°C until further used. The Quantitative PCR (qPCR) reaction was carried out using SYBR Green Supermix in a C1000 Touch Thermal Cycler CFX384 Real-Time System (Rio-Rad). The primers used for qPCR analysis are listed in **Methods S1**. ANOVA analysis of gene expression data was carried out with SAS software using the ΔΔCt method to determine gene expression changes (Livak and Schmittgen, 2001). QPCR analysis was carryout out using three biological replicates for H9 and H1 hESCs as specified in the figure panels.

### Immunofluorescence analysis

For immunofluorescence analysis, cells were grown on glass-bottom culture dishes coated with 3 μg/ml vitronectin and 1 μg/ml of Laminin 521. Cells were fixed either using 4% paraformaldehyde in PBS for 10 min, permeabilized with 0.5% Triton X-100 for 5 min and blocked in 3% BSA/PBS with 0.1% human IgG and 0.3% Triton X-100 for 1 hr. Cells were then incubated overnight with the primary antibody diluted in blocking buffer. The following primary antibodies were used: anti-KRT7 (SCB, 1:50), anti-KRT7 (CST, 1:500), rabbit anti-hCG (1:100), mouse anti-hCG (1:100), anti-YAP (1:200), anti-TFAP2C (1:400), anti-P63 (1:600), anti-GATA3 (1:500), anti-TEAD4 (1:250), anti-CDX2 (1:300), anti-VE-Cadherin (1:400), anti-HLA-G (1:300), anti-syncytin (1:50). Corresponding isotype controls (rabbit polyclonal IgG, rabbit XP IgG, mouse IgG1, and mouse IgG2a) were used at primary antibody concentrations. Alexa Fluor 488- or Alexa Fluor 647-conjugated secondary antibodies were used as secondary antibodies. Nuclei were stained with DAPI and all samples were imaged using a Zeiss LSM 710 or 880 laser scanning confocal microscope (Carl Zeiss, Germany).

### Confocal image analysis

Image analysis was conducted using an image processing algorithm created in MATLAB. First, the DAPI stain was isolated, binarized, and processed to accurately represent the number of cells in each image. The primary-antibody stain of interest was isolated and processed in the same manner. Only primary-antibody pixels that overlap DAPI pixels were considered for analysis, and the average intensities of those pixels were measured and correlated to the nearest nuclei. This was performed for one control image and multiple experimental images. Each cell in the experimental images was considered positively stained if the average intensity of that cell was greater than the average intensity of all of the cells in the control image. Statistical analysis was done using a two-tailed t-test evaluating percent positive cells from different treatment periods.

### RNA sequencing analysis using next generation sequencing

Total RNA was extracted with Trizol™ reagent using manufacturer’s protocol. RNA was purified using GeneJET RNA Purification Kit using manufacturer’s protocol. Isolated RNA samples were then used to evaluate genome-wide mRNA expression profiles using next generation RNA-sequencing, conducted at GENEWIZ, LLC. (South Plainfield, NJ, USA). RNA samples received at GENEWIZ were quantified using Qubit 2.0 Fluorometer (Life Technologies, Carlsbad, CA, USA) and RNA integrity was checked using Agilent TapeStation 4200 (Agilent Technologies, Palo Alto, CA, USA).

RNA sequencing libraries were prepared using the NEBNext Ultra RNA Library Prep Kit for Illumina following manufacturer’s instructions (NEB, Ipswich, MA, USA). Briefly, mRNAs were first enriched with Oligo(dT) beads. Enriched mRNAs were fragmented for 15 minutes at 94 °C. First strand and second strand cDNAs were subsequently synthesized. cDNA fragments were end repaired and adenylated at 3’ends, and universal adapters were ligated to cDNA fragments, followed by index addition and library enrichment by limited-cycle PCR. The sequencing libraries were validated on the Agilent TapeStation (Agilent Technologies, Palo Alto, CA, USA), and quantified by using Qubit 2.0 Fluorometer (Invitrogen, Carlsbad, CA) as well as by quantitative PCR (KAPA Biosystems, Wilmington, MA, USA).

The sequencing libraries were clustered on 4 lanes of a flowcell. After clustering, the flowcell was loaded on the Illumina HiSeq 4000 instrument according to manufacturer’s instructions. The samples were sequenced using a 2×150bp Paired End (PE) configuration. Image analysis and base calling were conducted by the HiSeq Control Software (HCS). Raw sequence data (.bcl files) generated from Illumina HiSeq was converted into fastq files and de-multiplexed using Illumina’s bcl2fastq 2.17 software. One mismatch was allowed for index sequence identification.

After investigating the quality of the raw data, sequence reads were trimmed to remove possible adapter sequences and nucleotides with poor quality using Trimmomatic v.0.36. The trimmed reads were mapped to the Homo sapiens GRCh38 reference genome available on ENSEMBL using the STAR aligner v.2.5.2b. The STAR aligner is a splice aligner that detects splice junctions and incorporates them to help align the entire read sequences. BAM files were generated as a result of this step. Unique gene hit counts were calculated by using feature Counts from the Subread package v.1.5.2. Only unique reads that fell within exon regions were counted.

### Analysis of gene expression profiles

After extraction of gene hit counts, the gene hit counts table was used for downstream differential expression analysis. Genome-wide RNA sequencing count data were processed and statistically assessed using the DESeq2 package (v1.22.2) in R Software (3.6.0) (The R Foundation, 2019). Count data were first filtered to include transcripts expressed above background, requiring the median across samples to be greater than the overall median signal intensity, as implemented in DESeq2. Count data were then normalized by median signal intensity using algorithms enabled within DESeq2, resulting in variance stabilized expression values (Love et al., 2014). These normalized values were used to carry out a principal component analysis (PCA) comparing data-reduced global expression signatures across samples. Principal components were calculated and visualized using the prcomp function in R (R-core, 2019). Heat maps were generated using Partek^®^ Genomics Suite Software (v7.18.0723) and gene-specific plots using GraphPad Prism Software (v8.2.0)), based on normalized expression values.

### Statistical and gene set enrichment analysis of genes differentially expressed between hTESCs and hTSCs

Genes that showed the greatest difference in expression between the hTESCs and hTSCs were identified using an analysis of variance analysis (ANOVA) comparing the normalized expression levels between these two groups. Genes showing the greatest difference in expression between the hTESCs and hTSCs were identified using the following statistical filters: (1) a false discovery rate-corrected q-value<0.05 (Storey, 2003), and (2) a fold change in expression (ratio of average across hTESCs over hTSCs samples) ≥ ± 1.5. To evaluate the biological role of these genes, a gene set enrichment analysis was carried out on the genes identified as significantly differentially expressed between groups. Specifically, all Gene Ontology (GO) gene sets (n=9996) from the Molecular Signature Database (MSigDB) (The Broad Institute, 2019) were queried for using the right-tailed Fisher’s Exact test, as enabled through the ‘platform for integrative analysis of omics data’ (PIANO) packing in R (Väremo et al., 2013). Gene sets were required to have an enrichment p-value<0.01 to be considered significant, consistent with previously published methods (Klaren et al., 2019; Rager et al., 2019). Genes that were identified at higher expression levels were evaluated separately from genes identified at significantly lower expression levels in hTESCs vs. hTSCs.

## Acknowledgments

This work was supported by NIH grants HD092741, HD093982 and NSF grant CBET 1706118

## Author contributions

Conceptualization: AM and BR; Investigation: AM, VK, JM; Formal Analysis: AM, VK, BR, CC, JR, RF; Data curation: AM, BR, JR; Resources: ASM; Writing – original draft: AM and BR; Writing – review and editing: AM, BR, VK, JR, RF; Visualization: AM, VK, CC, JR; Project Administration: BR; Funding Acquisition: BR and ASM

## Supplemental Information

Document S1.pdf (contains Supplemental Figures S1-S5, Methods S1)

Tables S1-S4.xlsx (contains Supplemental Tables S1-S4)

