## Supplementary material for "Two distinct trophectoderm lineage stem cells from human pluripotent stem cells": Document S1

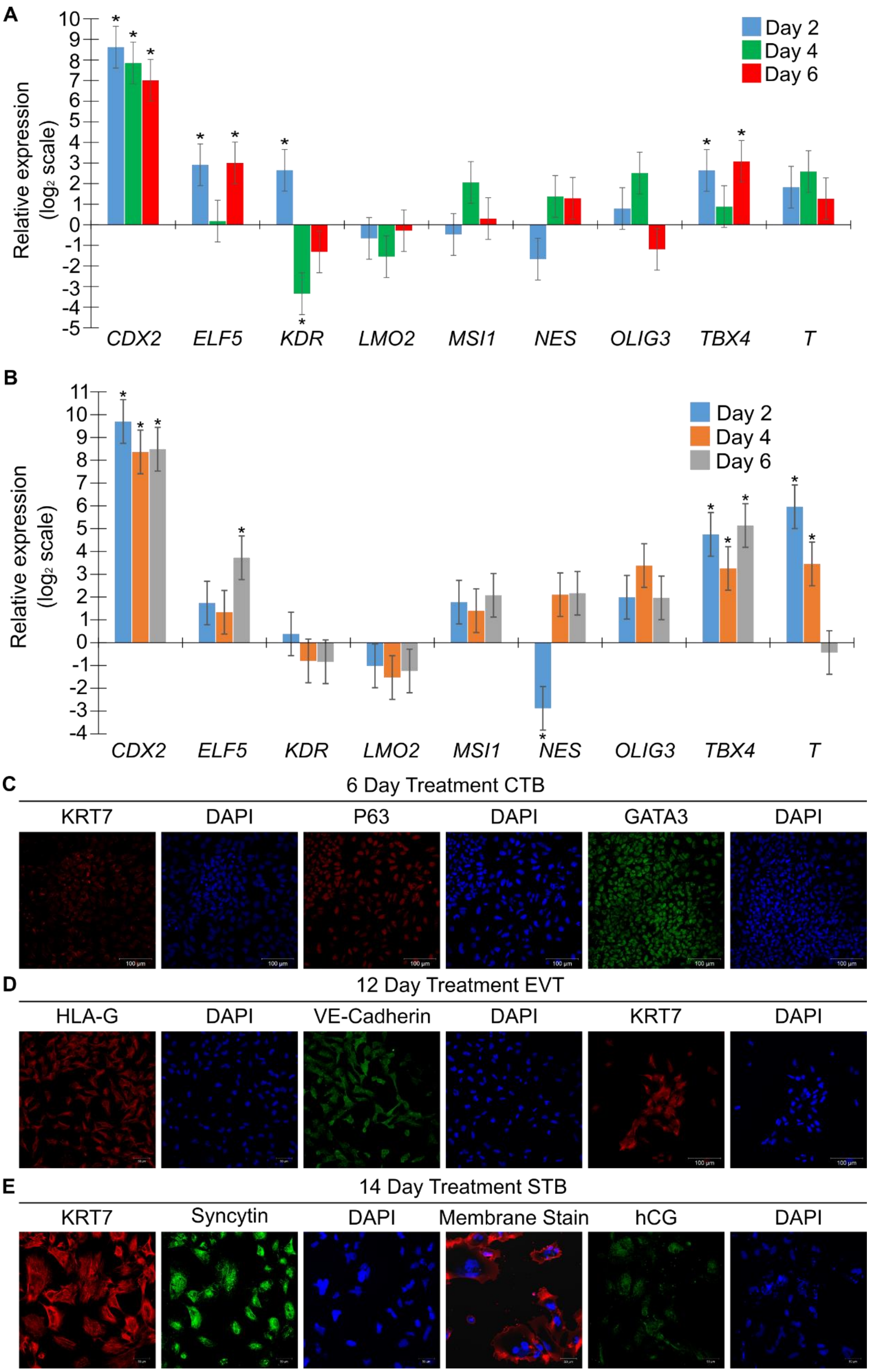

A
Relative expression (log2 scale)
Day 2
Day 4
Day 6
CDX2
ELF5
KDR
LMO2
MSI1
NES
OLIG3
TBX4
T
B
Relative expression (log2 scale)
Day 2
Day 4
Day 6
CDX2
ELF5
KDR
LMO2
MSI1
NES
OLIG3
TBX4
T
C
6 Day Treatment CTB
KRT7
DAPI
P63
DAPI
GATA3
DAPI
D
12 Day Treatment EVT
HLA-G
DAPI
VE-Cadherin
DAPI
KRT7
DAPI
E
14 Day Treatment STB
KRT7
Syncytin
DAPI
Membrane Stain
hCG
DAPI

**Figure S1: A chemically defined medium containing S1P enables differentiation of hESCs to CTB. Related to Figure 1.**

(A) Gene expression of *CDX2, ELF5, KDR, LMO2, MSH1, NES* (nestin), *OLIG3*, and *T* (brachyury) in H9 hESCs undergoing differentiation, compared to hESCs. Three biological replicates were used. (Error bars, S.E., *p<0.05).

(B) Gene expression of *CDX2, ELF5, KDR, LMO2, MSH1, NES* (nestin), *OLIG3*, and *T* (brachyury) in H1 hESCs undergoing differentiation compared to hESCs. Three biological replicates were used. (Error bars, S.E., *p<0.05).

(C) Immunostaining of KRT7, P63 and GATA3 in H1 hESCs at day 6 of initial treatment. Nuclei were stained with DAPI.

(D) Confocal images of EVT-like cells from 12-day treatment of H1 hESCs, staining for KRT7, HLA-G and VE-Cadherin. Nuclei were stained with DAPI.

(E) Confocal images of STB-like cells from 14-day treatment of H1 hESCs, staining for KRT7, syncytin and hCG. Nuclei were stained with DAPI.

Scale bars are 100μm for all images.

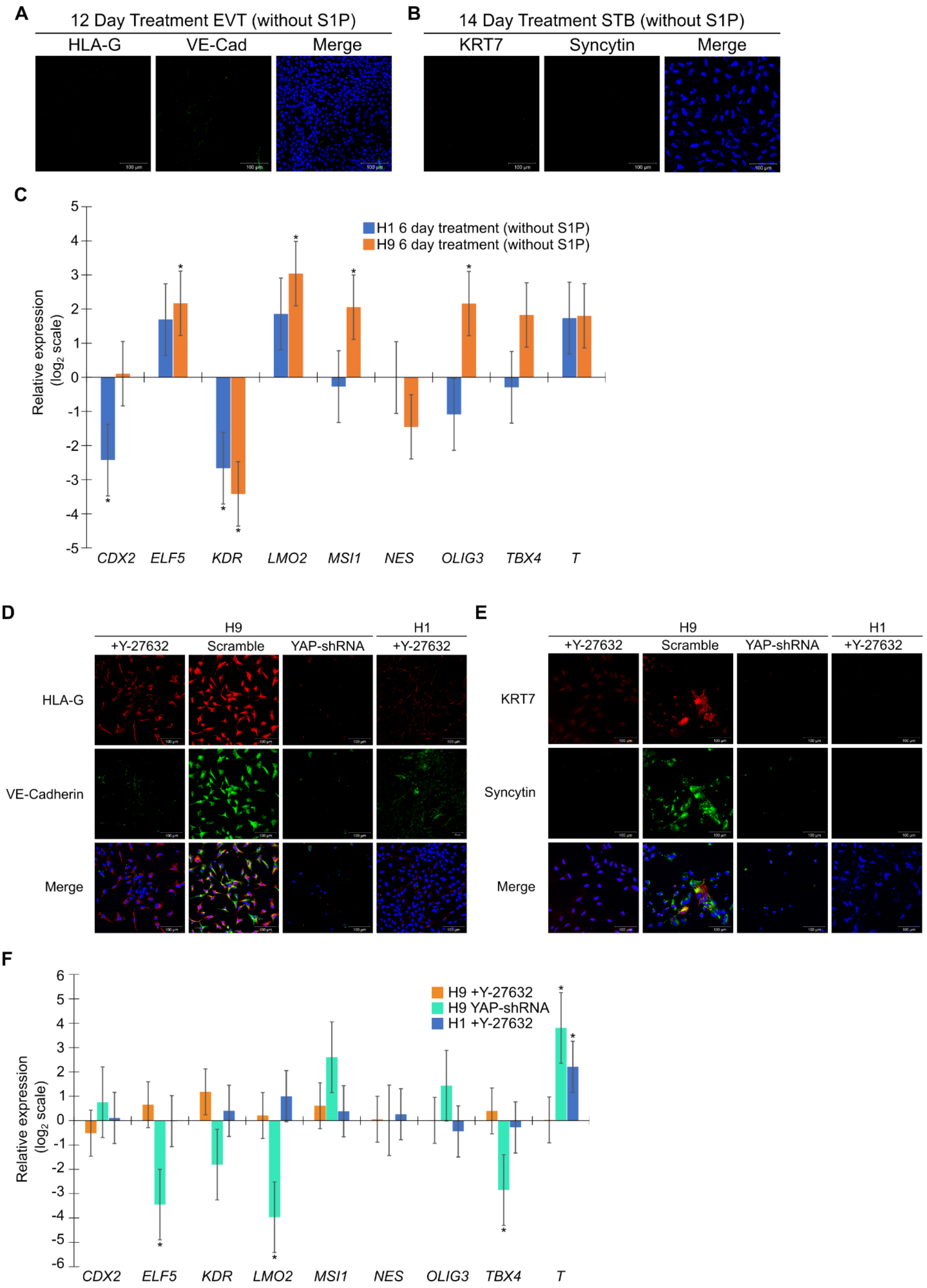
A
12 Day Treatment EVT (without S1P)
HLA-G
VE-Cad
Merge
B
14 Day Treatment STB (without S1P)
KRT7
Syncytin
Merge
C
H1 6 day treatment (without S1P)
H9 6 day treatment (without S1P)
Relative expression
(log2 scale)
5
4
3
2
1
0
-1
-2
-3
-4
-5
CDX2
ELF5
KDR
LMO2
MSI1
NES
OLIG3
TBX4
T
D
H9
H1
+Y-27632
Scramble
YAP-shRNA
+Y-27632
HLA-G
VE-Cadherin
Merge
E
H9
H1
+Y-27632
Scramble
YAP-shRNA
+Y-27632
KRT7
Syncytin
Merge
F
H9 +Y-27632
H9 YAP-shRNA
H1 +Y-27632
Relative expression
(log2 scale)
6
5
4
3
2
1
0
-1
-2
-3
-4
-5
-6
CDX2
ELF5
KDR
LMO2
MSI1
NES
OLIG3
TBX4
T

**Figure S2: S1P, Rho/Rock signaling, and YAP are necessary for differentiation of hESCs to trophoblast. Related to Figure 1.**

(A) Confocal images of cells from 12-day EVT treatment of H1 hESCs upon removal of S1P, staining for HLA-G and VE-Cadherin. Nuclei were stained with DAPI.

(B) Confocal images of cells from 14-day STB treatment of H1 hESCs upon removal of S1P staining for KRT7 and syncytin. Nuclei were stained with DAPI.

(C) Gene expression of *CDX2, ELF5, KDR, LMO2, MSH1, NES* (nestin), *OLIG3*, and *T* (brachyury) in H9 and H1 hESCs undergoing differentiation upon removal of S1P, compared to the 6-day time point in the presence of S1P. Three biological replicates were used. (Error bars are S.E., *$p<0.05$)

(D) Confocal images of cells from 12-day EVT treatment of H9 and H1 hESCs with the addition of ROCK inhibitor (+Y-27632), knockdown of YAP using an inducible shRNA (YAP-shRNA), or a scrambled shRNA control (scramble), staining for HLA-G and VE-Cadherin. Nuclei were stained with DAPI.

(E) Confocal images of cells from 14-day STB treatment of H9 and H1 hESCs with addition of ROCK inhibitor (+Y-27632), knockdown of YAP using an inducible shRNA (YAP-shRNA), or a scrambled shRNA control (scramble), staining for KRT7 and syncytin. Nuclei were stained with DAPI.

(F) Gene expression of *CDX2, ELF5, KDR, LMO2, MSH1, NES* (nestin), *OLIG3*, and *T* (brachyury) in H9 and H1 hESCs undergoing differentiation in the presence of S1P with the addition of ROCK inhibitor (+Y-27632) and knockdown of YAP using an inducible shRNA (YAP-shRNA), compared to the 6-day time point in the presence of S1P (for ROCK inhibitor) or scrambled shRNA control in the presence of S1P (for YAP knockdown). Three biological replicates were used. (Error bars are S.E., *$p<0.05$)

Scale bars are 100μm for all images.

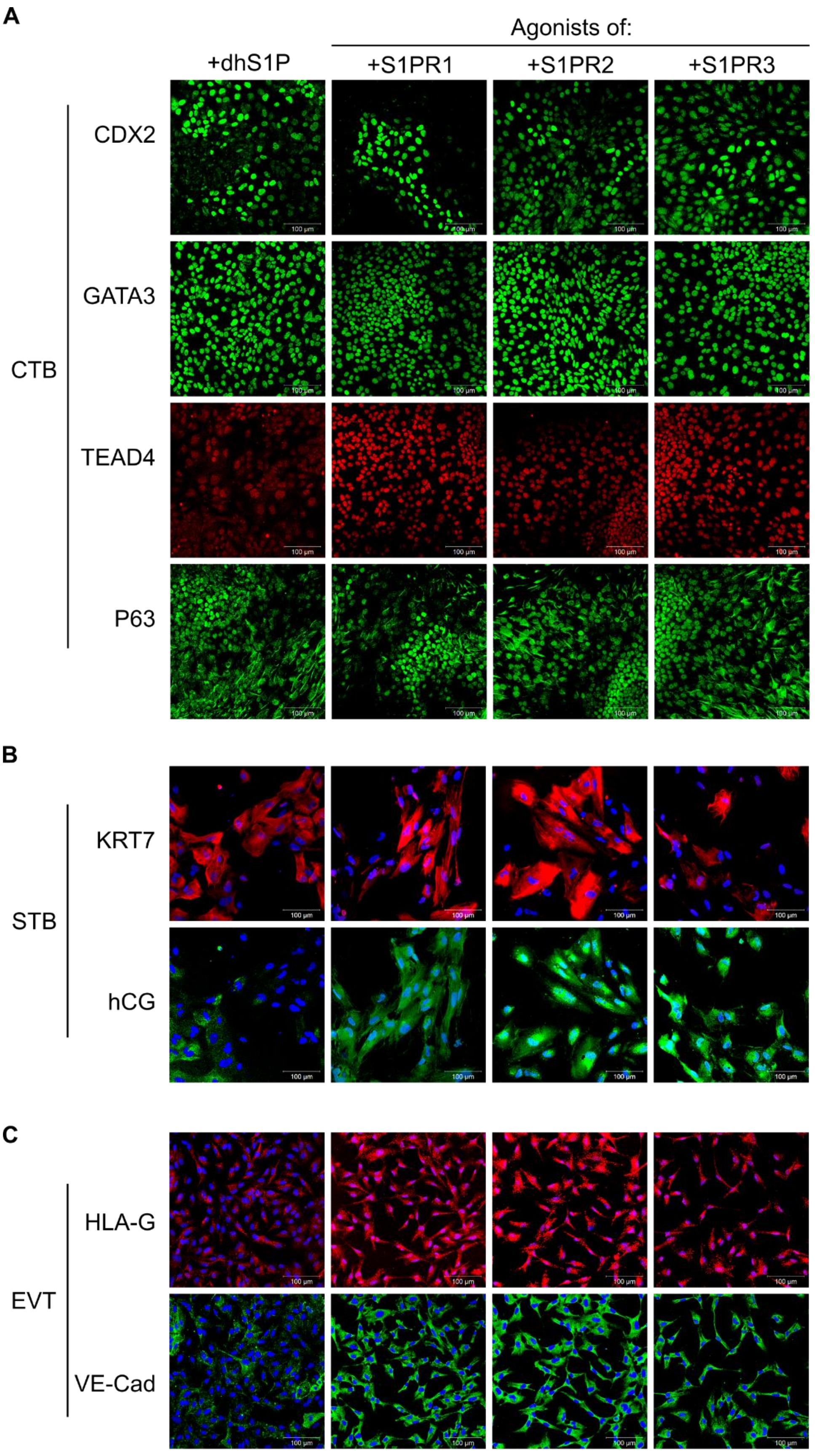

A
Agonists of:
+dhS1P
+S1PR1
+S1PR2
+S1PR3
CTB
CDX2
GATA3
TEAD4
P63
B
STB
KRT7
hCG
C
EVT
HLA-G
VE-Cad
100 μm

**Figure S3: S1P mediates its effects on trophoblast differentiation of hESCs through its receptors. Related to Figure 2.**

(A) Confocal images of CTB-like cells from 6-day treatment of H1 hESCs using D-erythro-dihydrospingosine-1-phosphate (dhS1P), CYM5442 (S1PR1 agonist), CYM5220 (S1PR2 agonist), and CYM5541 (S1PR3 agonist), staining for CDX2, GATA3, P63, and TEAD4. Nuclei were stained with DAPI.

(B) Confocal images of STB-like cells from 14-day treatment of H1 hESCs using dhS1P, CYM5442, CYM5520, and CYM5541 during the initial 6-day treatment, staining for KRT7 and hCG. Nuclei were stained with DAPI.

(C) Confocal images of EVT-like cells from 12-day treatment of H1 hESCs usingdhS1P, CYM5442, CYM5220, and CYM5541 during the initial 6-day treatment, staining for HLA-G and VE-Cadherin. Nuclei were stained with DAPI.

Scale bars are 100µm for all images.

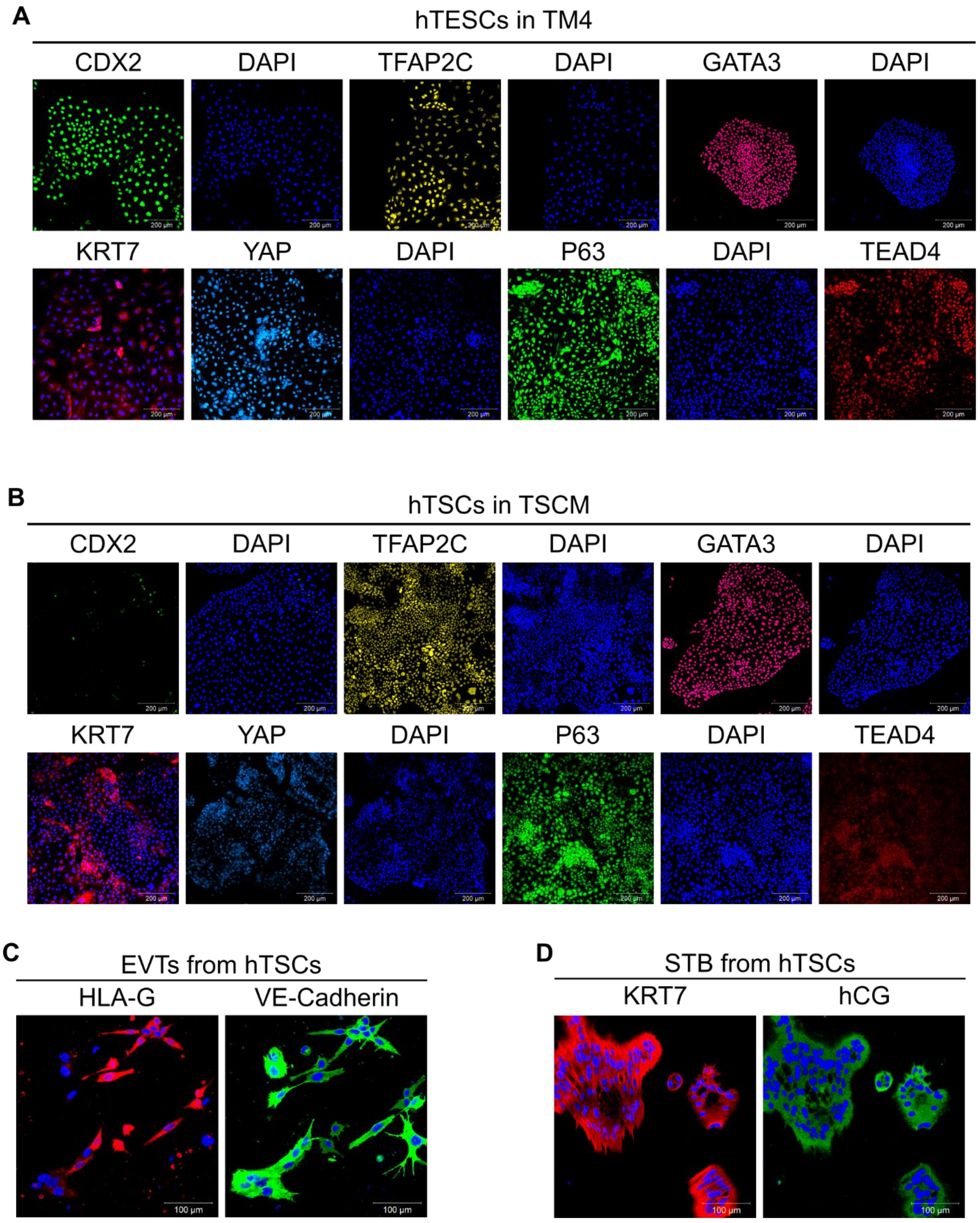
A
hTESCs in TM4
CDX2
DAPI
TFAP2C
DAPI
GATA3
DAPI
KRT7
YAP
DAPI
P63
DAPI
TEAD4
B
hTSCs in TSCM
CDX2
DAPI
TFAP2C
DAPI
GATA3
DAPI
KRT7
YAP
DAPI
P63
DAPI
TEAD4
C
EVTs from hTSCs
HLA-G
VE-Cadherin
D
STB from hTSCs
KRT7
hCG

**Figure S4: Optimizing timing of hESC differentiation enables derivation of CDX2+ hTESCs and P63+ hTSCs. Related to Figure 3.**

(A) Confocal images of H1 hESC-derived hTESCs in TM4, staining for CDX2, TFAP2C and GATA3, YAP, TEAD4, and P63. Nuclei were stained with DAPI. Scale bars are 200 µm.

(B) Confocal images of H1 hESC-derived hTSCs in TSCM, staining for CDX2, TFAP2C and GATA3, YAP, TEAD4, and P63. Nuclei were stained with DAPI. Scale bars are 200 µm.

(C) Confocal images of EVTs from H1 hESC-derived hTSCs, staining for HLA-G and VE-Cadherin. Nuclei were stained with DAPI. Scale bars are 200 µm.

(D) Confocal images of STB from H1 hESC-derived hTSCs staining for hCG and KRT7. Nuclei were stained with DAPI. Scale bars are 100µm

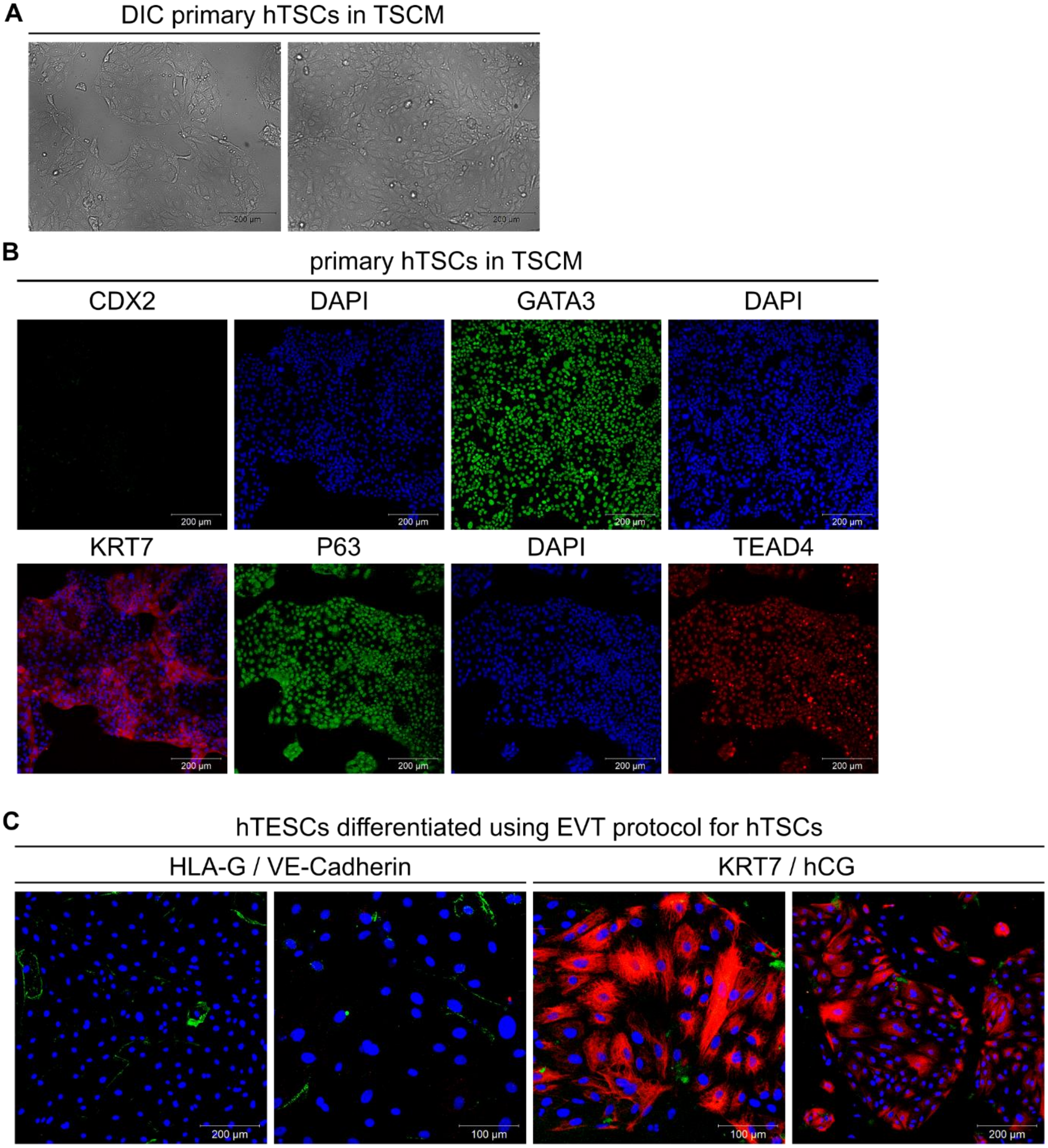
A
DIC primary hTSCs in TSCM
200 µm
200 µm
B
primary hTSCs in TSCM
CDX2
DAPI
GATA3
DAPI
KRT7
P63
DAPI
TEAD4
200 µm
C
hTESCs differentiated using EVT protocol for hTSCs
HLA-G / VE-Cadherin
KRT7 / hCG
200 µm
100 µm
100 µm
200 µm

**Figure S5: Related to Figure 3 and Figure 5.**

Primary hTSCs cultured in TSCM

(A) DIC images of primary CT29 hTSCs cells in TSCM

(B) Confocal images of primary CT29 hTSCs staining for CDX2, GATA3, TEAD4, and P63 grown in TSCM. Nuclei were stained with DAPI. Scale bars are 200µm for all images.

hTESCs do not differentiate to EVTs using the same protocol as that used for hTSCs

(C) Confocal images of cells from H9-derived hTESCs, simultaneously staining for HLA-G (red) and VE-Cadherin (green), or hCG (green) and KRT7 (red). Staining for HLA-G is not observed. Minimal staining for VE-Cadherin and hCG is observed. Nuclei were stained with DAPI. Scale bars are 200µm and 100µm.

### Methods S1: List of primers used for quantitative PCR analysis

| Gene | Primer | Sequence |
|---|---|---|
| ***CDX2*** | Forward | GGC AGC CAA GTG AAA ACC AG |
| ***CDX2*** | Reverse | GGT GAT GTA GCG ACT GTA GTG AA |
| ***ELF5*** | Forward | GCT GCG ACC AGT ACA AGT TG |
| ***ELF5*** | Reverse | CTG CCT CGA CGA ACT CCT C |
| ***GAPDH*** | Forward | CTC CAC GAC GTA CTC AGC G |
| ***GAPDH*** | Reverse | TGT TGC CAT CAA TGA CCC CTT |
| ***KDR*** | Forward | GGC CCA ATA ATC AGA GTG GCA |
| ***KDR*** | Reverse | CCA GTG TCA TTT CCG ATC ACT TT |
| ***LMO2*** | Forward | GGC CAT CGA AAG GAA GAG CC |
| ***LMO2*** | Reverse | GGC CCA GTT TGT AGT AGA GGC |
| ***MSI1*** | Forward | TAA AGT GCT GGC GCA ATC G |
| ***MSI1*** | Reverse | TCT TCT TCG TTC GAG TCA CCA |
| ***NES*** | Forward | CTG CTA CCC TTG AGA CAC CTG |
| ***NES*** | Reverse | GGG CTC TGA TCT CTG CAT CTA C |
| ***OLIG3*** | Forward | AGC CGT CTC AAC TCG GTC T |
| ***OLIG3*** | Reverse | CAT GGC TAG GTT CAG GTC GTG |
| ***T*** | Forward | CTG GGT ACT CCC AAT GGG G |
| ***T*** | Reverse | GGT TGG AGA ATT GTT CCG ATG A |
| ***TBX4*** | Forward | TGT TCC CCA GCT ACA AGG TAA |
| ***TBX4*** | Reverse | GCA GGG ACA ATG TCA ATC AGC |
